# Identification of the key role of white matter alteration in the pathogenesis of Huntington’s Disease

**DOI:** 10.1101/2021.06.21.449242

**Authors:** Jean-Baptiste Pérot, Marina Célestine, Marco Palombo, Marc Dhenain, Sandrine Humbert, Emmanuel Brouillet, Julien Flament

## Abstract

Pathogenesis of the inherited neurodegenerative disorder Huntington’s Disease (HD) is complex and progressive, with a long presymptomatic phase in which subtle changes occur in the brain of gene carriers up to 15 years before the onset of symptoms. Thus, there is a need of early, functional biomarker to better understand disease progression and to evaluate treatment efficacy far from onset. In particular, recent studies have shown that white matter may be affected early in HD. In this study, we scanned longitudinally Ki140CAG mice with structural MRI, Diffusion Tensor Imaging (DTI), Chemical Exchange Saturation Transfer of glutamate (gluCEST) and Magnetization Transfer (MT) imaging, in order to assess white matter integrity over the life of this very progressive mouse model. Our results show early defects of diffusion properties in the anterior part of the corpus callosum, preceding gluCEST defects in the same region (−10.8% at 8 months, −19% at 12 months) that spread to adjacent regions. At 12 months, frontal (−7.3%) and piriform (−16.7%) cortices showed reduced gluCEST, as well as the pallidum (−21.0%). MT imaging showed reduced signal in the septum (−21.7%) at 12 months. Cortical and striatal atrophy then appear at 18 months. Vulnerability of the striatum and motor cortex, combined with alterations of anterior corpus callosum, seems to point out the pivotal role of white matter, in the pathogenesis of HD and the pertinence of gluCEST and DTI as biomarkers in HD.

**Highlights:**

– A knock-in mouse model of Huntington’s disease is longitudinally characterized
– A multimodal MRI protocol is performed to identify biomarkers of the disease
– The white matter plays a pivotal role in the pathogenesis of the disease
– The cortico-striatal pathway seems particularly vulnerable in Huntington’s disease

## 1. Introduction

Huntington’s disease (HD) is an inherited neurodegenerative disease characterized by a triad of symptoms. While the disease is well known for its most prominent motor symptoms (chorea, dyskinesia and dystonia), many non-motor manifestations also occur early including olfactory deficits, sleep disorders, psychiatric signs such as depression, perturbation of emotivity and irritability. Cognitive deficits are also observed including loss of mental flexibility/adaptation, and defective procedural memory ^1^. Despite tremendous efforts made, there is currently no therapy to cure the disease or to slow its progression, resulting in a fatal outcome 10 to 15 years after the onset of symptoms in young individuals. The disease is caused by a mutation consisting of an abnormal expansion of CAG repeats in the exon 1 of the HTT gene coding the protein huntingtin (htt) ^2^. This leads to an expansion of a poly-glutamine tract in the N-terminal part of the protein. Mutant htt (m-htt) is toxic to brain cells through combination of gain and loss of function mechanisms ^3^.

Non-motor manifestations in m-HTT gene carriers are among the earliest to occur and are very disabling features in young individuals. Several studies suggested that they could occur many years before the onset of the disease ^4,5^. However, these manifestations can be highly variable from one patient to the other and their use as clinical readouts for future treatment evaluation is difficult, as it requires important cohort sizes. Similarly, the intra- and inter-subject variability of motor symptoms, although lower than that of non-motor signs, also limits their use to monitor disease progression ^6,7^.

In addition to symptoms, neuroimaging methods likely represent a crucial non-invasive approach to characterize in a quantitative manner the progression of the disease at the individual level. The preferential atrophy and dysfunction of the striatum first evidenced as the most striking neuropathological finding in post-mortem HD brain samples ^8,9^ was confirmed more than two decades ago by founding studies using CT, MRI and PET scanners even in patients at early stages of the disease ^10,11^. Recent work showed that the atrophy of the striatum as measured by MRI is currently the best biomarker of disease progression in HD gene carriers ^12,13^. However, despite a good correlation between striatal atrophy and disease severity ^14^, atrophy provides only limited information about HD pathogenesis and is probably the long-term consequence of subtle biological modifications that occur many years before macroscopic alterations. In addition, numerous studies suggested that volume alterations could occur simultaneously in several other brain structures such as cortical and subcortical regions ^15,16^ as well as white matter (WM) ^17,18^. Therefore, focusing only on striatal atrophy is too limiting to capture the extent of different biological processes involved in HD. Thus, there is a need to find more predictive, functional and earlier biomarkers for a better understanding of disease pathogenesis and to monitor its progression. Moreover, in the perspective of future therapeutic interventions aiming at slowing down disease progression before onset of disabling symptoms, such biomarkers would be extremely useful to evaluate drug engagement and biological efficacy.

Interestingly, recent studies pointed out early alterations of WM and fiber tracks in HD concomitantly with atrophy measured in gray matter (GM) ^17,19^. Furthermore, longitudinal studies performed in large cohorts of HD gene carriers such as TRACK-HD identified a good correlation between atrophy measured in frontal and posterior WM and disease progression in presymptomatic HD patients ^14,20^. Early WM changes have also been reported in animal models of HD where anomalies of WM tracks were found using MRI ^21,22^. In addition, the presence of m-htt in oligodendrocytes of knock-in mice has been shown to lead to abnormal myelination and apparition of neurological symptoms ^23,24^. Although WM changes have been considered as a consequence of GM loss for many years, numerous studies provided strong evidence of the major role of WM alterations in HD physiopathology, independently of neuronal cell loss ^25–27^. Therefore, WM modifications might be one of the early hallmarks of the disease and could constitute a key biomarker, especially during the presymptomatic phase of the disease.

Further investigations are needed to characterize WM changes induced by HD pathological processes and neuroimaging methods can provide valuable tools to address this question. MRI can be used to assess non-invasively several aspects of the pathology, going from neuro-metabolic modifications to macrostructural changes.

Numerous studies have used Diffusion Tensor Imaging (DTI) to analyze WM modifications ^28^. Quantitative indexes such as fractional anisotropy (FA), mean diffusivity (MD), radial diffusivity (RD, diffusivity perpendicular to axonal fibers) and axial diffusivity (AD, diffusivity along axonal fibers) can be derived to assess microstructural properties of WM ^29^. FA describes the degree of anisotropy within a voxel so a decrease of FA value represents a loss of tissue organization that can be associated with alteration of cellular integrity ^30^. AD and RD are usually considered to be increased in pathological conditions due respectively to axonal degeneration and demyelinating processes ^31^. However, several studies have shown inconsistent results in the context of HD so interpretation of these variations remains ambiguous ^27,32,33^. Such heterogeneity may be explained by the variability of symptoms within HD patients or due to different characteristics of animal models.

Complementarily to diffusion MRI, magnetization transfer (MT) imaging can capture changes of macromolecular content of tissues with a good spatial resolution, especially myelin content ^34^. The MT contrast relies on decrease of water signal intensity due to exchange of magnetization between fast exchanging protons bound to macromolecules and free water after an off-resonance radio frequency (RF) pulse ^35^. This method has been shown to be sensitive to myelin concentration with a good specificity and several studies used it to explore both WM and GM changes in HD patients ^36,37^.

In vivo ^1^H Magnetic Resonance Spectroscopy (^1^H-MRS) can monitor alterations of metabolic profiles, providing key biological insights of pathological pathways involved in different cell types ^38,39^. Previous studies demonstrated metabolic changes that occur in the brain of either human HD patients or transgenic mouse models of HD ^40–42^. In particular, decreased concentrations of N-acetyl-aspartate (NAA) and glutamate (Glu), two metabolites mainly located in neurons, were measured in HD patients and animal models ^43^. Other metabolite concentrations such as glutamine (Gln), myo-inositol (Ins) and choline (Cho) have been reported to be altered in HD mouse models, even if variations were not consistently reported depending on the model ^44,45^. However, in spite of the rich biological information provided by ^1^H-MRS, this method suffers from its inherently low sensitivity. Measurement of metabolic profiles within small and thin structures such as the corpus callosum would be very difficult without strong contamination by surrounding structures.

Recently, CEST (Chemical Exchange Saturation Transfer) has been proposed to address this limitation by using indirect detection of dilute molecules through their exchange of labile protons with the bulk water ^46–48^. Exchangeable protons such as amine (-NH_2_) or amide (-NH) exhibit a resonance frequency that is shifted from the water resonance frequency so they can be selectively saturated using RF pulse. The decrease of water signal due to magnetization exchange being directly correlated to the concentration of exchanging protons, it is possible to map the spatial distribution of the molecule of interest. Several studies already demonstrated the potential of CEST imaging to map glutamate with a good spatial resolution in both rodent and human brains ^49–51^, including in the context of HD ^52^.

The Ki140CAG mouse model ^53^ is a knock-in model of HD containing 140 CAG repeats inserted in the murine huntingtin gene. It is characterized by the slowly progressive appearance of the symptoms and is considered to mimic more closely clinical form of HD than severe and rapid mouse models ^53,54^. These mice present early locomotor abnormalities and huntingtin aggregates in the striatum as early as 4 months of age ^53^. Homozygous mice for the HTT gene present early impairments of motor functions, such as hypokinesia at 4 months of age and reduced performances in rotarod test ^55^. In a recent study performed on heterozygous and homozygous Ki140CAG mice, we showed a progression of the severity of locomotor activity defects with genotype when compared to age-matched wild type littermates ^52^. Previously, we detected early brain changes using neuroimaging in this model. Structural MRI revealed significant striatal atrophy at 12 months of age in small cohorts of homozygous mice ^52^. ^1^H-MRS in the same mice showed metabolic modifications including reduced levels of taurine and NAA in the striatum, in both homozygous and heterozygous Ki140CAG mice. Concentration of Glu was also found to be reduced while Gln was increased. These alterations were associated with a decrease of gluCEST signal in the striatum and piriform cortex of homozygous mice. Surprisingly, the most affected structure was the corpus callosum in both heterozygous and homozygous mice, suggesting that this structure was very early affected in this mouse model. However, whereas all changes we observed tended to be stronger in homozygous than in heterozygous Ki140CAG mice, they were obtained only at one age, so that the progression of the disease in this model remained to be investigated.

In the present study, we developed a multimodal MRI protocol combining anatomical, diffusion weighted, gluCEST and MT imaging in order to monitor longitudinally Ki140CAG mice and their wild type (WT) littermates. We detected pointed toward early changes affecting the function and tissue organization of different extra-striatal brain regions providing new insights on the progression of brain perturbations in this slowly progressive model of HD that could be reminiscent of those occurring in m-HTT human gene carriers.

## 2. Material and Methods

### 2.1 Ki140CAG and wild type littermate mice

Knock-in mice express chimeric mouse/human exon 1 containing 140 CAG repeats inserted in the murine HTT gene (Ki140CAG) ^53^. Ki140CAG mice colony was maintained by breeding heterozygous Ki140CAG males and WT females. Heterozygous mice (n = 11 males, Ki140CAG) were compared to their relative age-matched littermates (n = 12 males, WT). Mice were housed in a temperature-controlled room maintained on a 12 hours light/dark cycle. Food and water were available ad libitum. All animal studies were conducted according to the French regulation (EU Directive 2010/63/EU – French Act Rural Code R 214-87 to 126). The animal facility was approved by veterinarian inspectors (authorization n° A 92-032-02) and complies with Standards for Humane Care and Use of Laboratory Animals of the Office of Laboratory Animal Welfare (OLAW – n°#A5826-01). All procedures received approval from the ethical committee (APAFIS #21335-201907031642584 v2).

### 2.2 MRI Protocol

Animals were scanned longitudinally (2.5, 5, 8, 12 and 18 months of age) on a horizontal 11.7 T Bruker scanner (Bruker, Ettlingen, Germany). Mice were first anesthetized using 3% isoflurane in a 1:1 gas mixture of air/O_2_ and positioned in a dedicated stereotaxic frame with mouth and ear bars to prevent any movements during MR acquisitions. Mice temperature was monitored with a rectal probe and maintained at 37°C with regulated water flow. Respiratory rate was continuously monitored using PC SAM software (Small Animal Instruments, Inc., Stony Brook, NY, USA) during scanning. The isoflurane level was adjusted around 1.5% to keep the respiratory rate in the range of 60 to 80 per minute. A quadrature cryoprobe (Bruker, Ettlingen, Germany) was used for radiofrequency transmission and reception.

High-resolution anatomical images (Turbo Spin Echo sequence, TE/TR = 5/10000 ms, Turbo factor = 10, effective TE = 45 ms, in-plane resolution = 100 × 100 μm^2^, 200 μm slice thickness, 100 slices) were used for structures volumetry.

Three gluCEST images centered on the mid-striatum were acquired with a 2D fast spin-echo sequence preceded by a frequency-selective continuous wave saturation pulse (TE/TR = 6/5000 ms, 10 echoes and effective TE = 30 ms, in-plane resolution = 150 × 150 μm^2^, 0.6 mm slice thickness). The MAPSHIM routine was applied in a voxel encompassing the slices of interest in order to reach a good shim on gluCEST images. GluCEST images were acquired with a saturation pulse applied during T_sat_ = 1 s, composed by 10 broad pulse of 100 ms, with 20 μs inter-delay and an amplitude B_1_ = 5 μT. The frequency of the saturation pulse Δω was applied in a range from −5 ppm to 5 ppm with a step of 0.5 ppm. The WASSR method 56 was used to correct for B_0_ inhomogeneities (B1 = 0.2 μT, Δω in a range from −1 ppm to 1 ppm with a step of 0.1 ppm). The same sequence than gluCEST was used for MT images acquisition with optimized parameters to assess macromolecular compounds (Δω = ±16 ppm, B_1_ = 10 μT, T_sat_ = 800 ms). MT images were not acquired during the first time-point (2.5 months).

The diffusion-weighted MRI data were acquired using Echo Planar Imaging (EPI) sequence (TE/TR = 30/3200 ms, in-plane resolution = 112 × 112 μm^2^, 500 μm slice thickness, 10 slices, b-value = 1000 s/mm^2^, 30 directions).

### 2.3 Data processing and statistical analyses

Images acquired using each of the MRI modalities were co-registered and automatically segmented using an in-house python library (Sammba-MRI ^57^, **Fig. 1**). In a first step, anatomical images were co-registered to generate an average of all anatomical images, called the study template. This template was segmented using an atlas composed by 34 regions (**Fig. 1**) derived from the Allen mouse brain atlas ^58^. Then, transformations to match the atlas to the study template and to match the study template to all high-resolution anatomical images were calculated. Finally, transformations were computed to project the atlas onto the images acquired using each of the MRI modalities. Outputs of the Sammba-MRI pipeline are the volume of each of the 34 structures and mean gluCEST contrast, MT contrast and FA, RD and AD within each structure for each animal.

**Figure 1:**
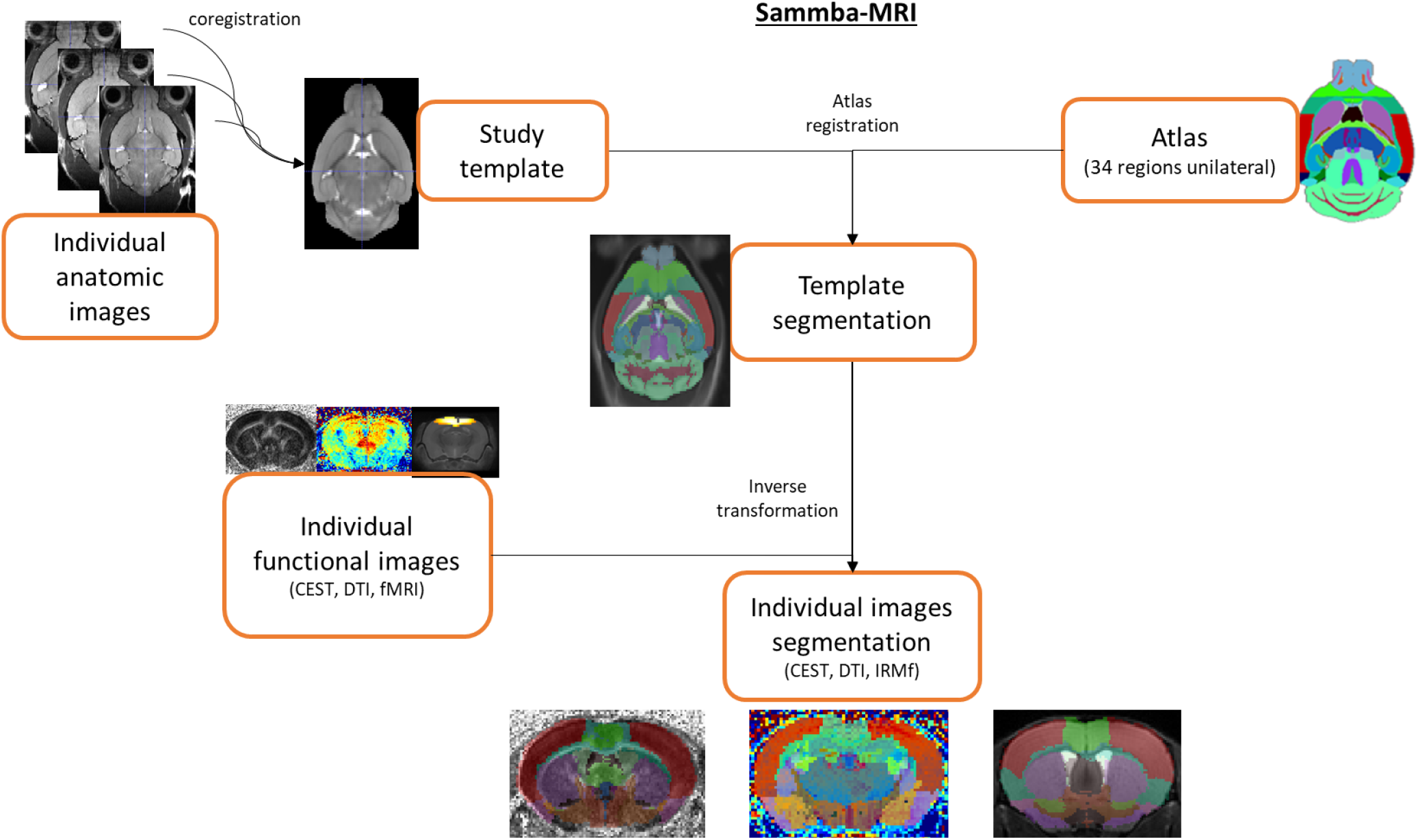
Automated co-registration and segmentation pipeline using in-house python library Sammba-MRI.

CEST images were processed pixel-by-pixel and analyzed using in-house programs developed on MATLAB software (MathWorks Inc., Natick, MA) used to generate Z-spectra by plotting the longitudinal magnetization as a function of saturation frequency. WASSR method was used to generate absolute B_0_ map by finding the actual frequency within each voxel. Z-spectrum in each voxel was interpolated using a cubic spline and B_0_ map was used to correct for B_0_ inhomogeneities. The specific glutamate contribution was isolated using Asymmetrical Magnetization Transfer Ratio (MTRasym) 59 and was calculated as follows: MTRasym(Δω) = 100 × (M_sat_(−Δω) - M_sat_(+Δω)) / M_sat_(−5 ppm) where M_sat_(±Δω) is the magnetization acquired with saturation pulse applied at ‘+’ or ‘-’ Δω ppm. GluCEST images were calculated with Δω centered at ± 3 ppm.

FA, AD and RD maps were generated using ParaVision 6.1 software (Bruker, Ettlingen, Germany). FA images underwent the Tract-Based Spatial Statistics (TBSS) pipeline ^60^ using FSL software ^61^. For that, FA maps were first co-registered to the study template. Then, the mean FA map over the whole animal cohort was generated and the skeleton template of WM tracks was extracted. Individual skeleton from each animal was calculated using the same process and then projected to skeleton template to have a good co-registration of WM tracks.

Variation of each index *Ind* (with *Ind* being either structure volume, gluCEST contrast or MT contrast) between WT littermates and Ki140CAG mice was calculated in each brain region as follows: Variation = 100 × (*Ind*(WT) - *Ind*(Ki140CAG)) / *Ind*(WT).

Statistical analyses were performed using GraphPad Prism 8.0 (GraphPad Software, San Diego, California, USA). The Shapiro-Wilk test was used to test the data for normality and no deviation from normality was observed for any data. One-way ANOVA with repeated measures was used for statistical analysis. The significant threshold was set to 0.05. ANOVA was followed by a Bonferroni *post-hoc* test to determine individual significant differences between groups. Sample size to compare HD and WT mice that would have been needed for detection of significant change was estimated assuming a significance level of 5%, a power of 80% and two-sided tests (BiostaTGV module, https://biostatgv.sentiweb.fr/?module=etudes/sujets#). The standard deviation used for this estimation was the square root of the pooled variance from each group.

### 2.4 Graph Theory

Normalized projection data from the Allen Connectivity Atlas (Oh et al.^62^) was processed using graph theory. This method has been widely used for representation of functional networks as measured by resting-state functional MRI. A threshold was applied at 1 of normalized projection strength to remove irrelevant projections. Thresholded projection matrix was then loaded in Gephi 0.9.2 and a Force Atlas spatialization algorithm was applied. This algorithm iteratively forms clusters of regions based on their connectivity. Correspondence between subregions from connectivity atlas and large structures used in MRI atlas was established based on the Allen Mouse Brain Atlas ^58^ (http://atlas.brain-map.org/).

### 2.5 Tractography

Acquired diffusion-weighted MRI data were reconstructed with the generalized q-sampling imaging (GQI) method ^63^, which models diffusion patterns in each voxel with an orientation distribution function (ODF) that can detect simultaneous diffusion in multiple directions, using the DSI Studio software (http://dsi-studio.labsolver.org). For tractography, the generalized deterministic tracking algorithm implemented in DSI Studio ^64^ was used. Whole brain tractography was first performed to assess overall data quality and to decide appropriate values for global parameters (e.g. the anisotropy threshold used as a stopping criterion in tractography) by tracking streamlines with a whole-brain seed. Then, locally-constrained tractography was used to isolate the tract connecting the cortex to the striatum, namely the cortico-striatal tract, and those connecting the two hemispheres, namely inter-hemispheric tract, by making use of ROI-based Boolean operations. For the cortico-striatal tract, ROIs were defined by specifying the volume of the motor cortex from which streamlines must start and the volume of the striatum where streamlines must terminate. For the inter-hemispheric tract, ROIs were defined by specifying the volume of the left end of the corpus callosum from which streamlines must start and the volume of the right end of the corpus callosum where streamlines must terminate, with an additional ROI in the middle of the corpus callosum that streamlines must cross. A threshold on the FA at 0.15, maximum streamline length 20 mm; random maximum angle; trilinear interpolation were used as global parameters. For whole brain tractography 100,000 streamlines were generated with random seeds in the whole brain; these streamlines where then filtered in the locally-constrained tractography step, where only the streamlines satisfying the ROI-based Boolean conditions were retained to define the cortico-striatal tract and the inter-hemispheric tract.

## 3. Results

### 3.1 *Long-term and evolving cortico-striatal atrophy in* Ki140CAG mice

The mean volume of each of the 34 brain regions was calculated at each time-point for WT littermates and Ki140CAG mice. Volume variation between mice groups was calculated for all brain structures and reported in a variation map in order to highlight hypertrophied (shades of pink) and atrophied (shades of green) structures (**Fig. 2**, variation expressed in percentage referred to WT littermates). At 18 months, significant atrophy of the striatum (−4.3%, p<0.05), frontal cortex (−5.3%, p<0.05) and motor cortex (−3.3%, p<0.05) were measured in Ki140CAG mice (**Fig. 2**, last column). At earlier time points, we saw a clear trend towards cerebral atrophy particularly in cortical regions in association with dilation of ventricles. Although volume variations were not significant, the atrophy trend seemed increased with aging.

**Figure 2:**
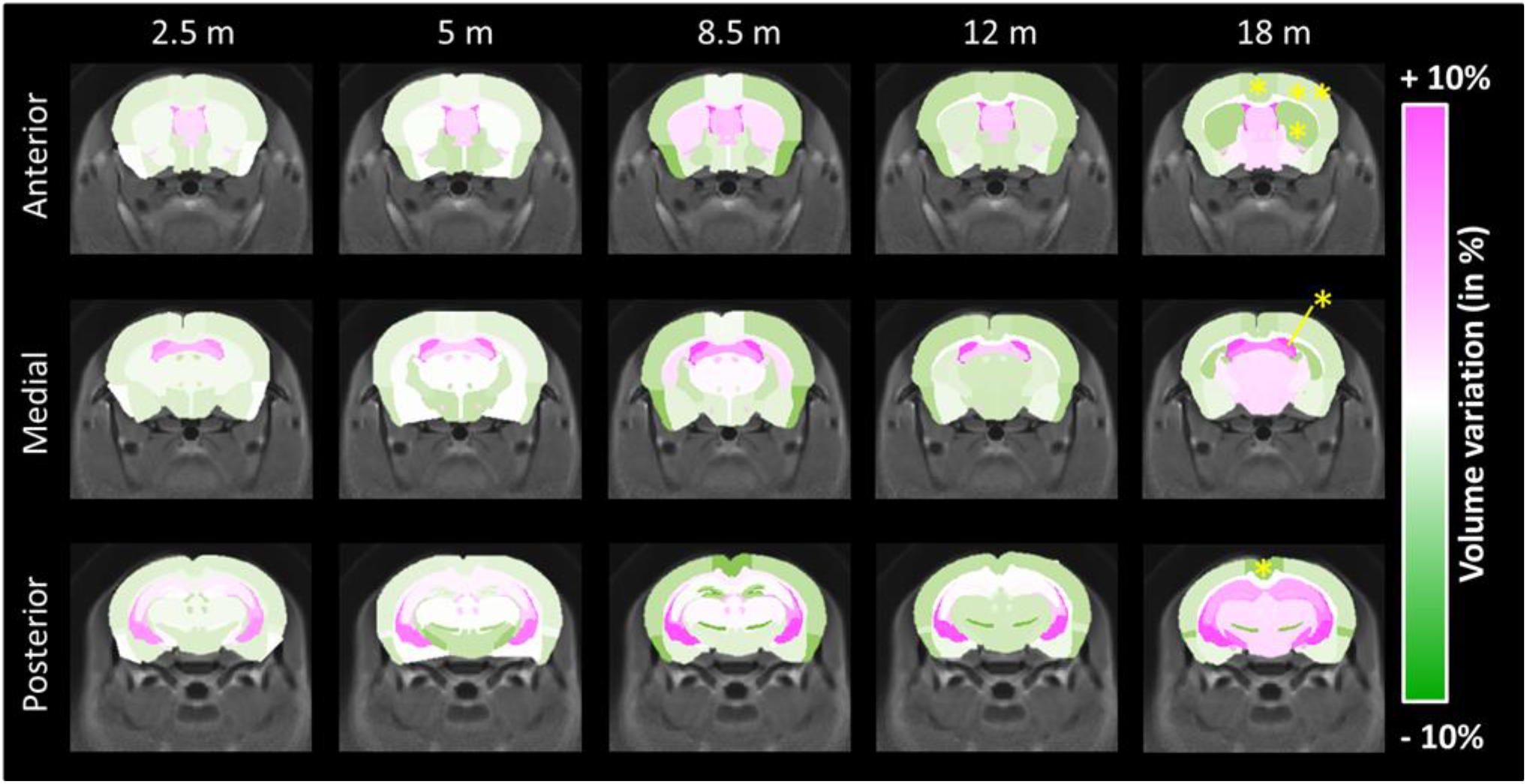
Volume variation maps between WT and Ki140CAG mice, at each time-point, on three slices of the mouse brain. Hypertrophied structures are reported in shades of pink and atrophied structures in shades of green. Yellow stars represent significant atrophy (RM-ANOVA + Bonferroni, p < 0.05).

### 3.2 Early deficiencies of cortico-striatal pathway revealed by metabolic imaging

Variations of mean gluCEST contrast (**Fig. 3**, increase of gluCEST contrast in shades of red, decrease in shades of blue) measured in each brain region at each time-point were calculated between Ki140CAG mice and WT littermates and reported on a variation map. The most affected structure in Ki140CAG mice was the corpus callosum (CC) where a significant decrease of gluCEST contrast was measured at 8 months (−10.8%, p < 0.05, **Fig. 3**). Significant decreases of gluCEST contrast were also measured in CC (−19%, p < 0.01), in the frontal (−7.3%, p < 0.05) and piriform (−16.7%, p < 0.05) cortices and in the pallidum (−21.0%, p < 0.05) of Ki140CAG mice at 12 months, confirming our previous results acquired in this mouse model ^52^. A non-significant decrease of gluCEST contrast (−12%, p = 0.12) was measured in the striatum of 12-months old mice, as already reported in heterozygous Ki140CAG mice.

**Figure 3:**
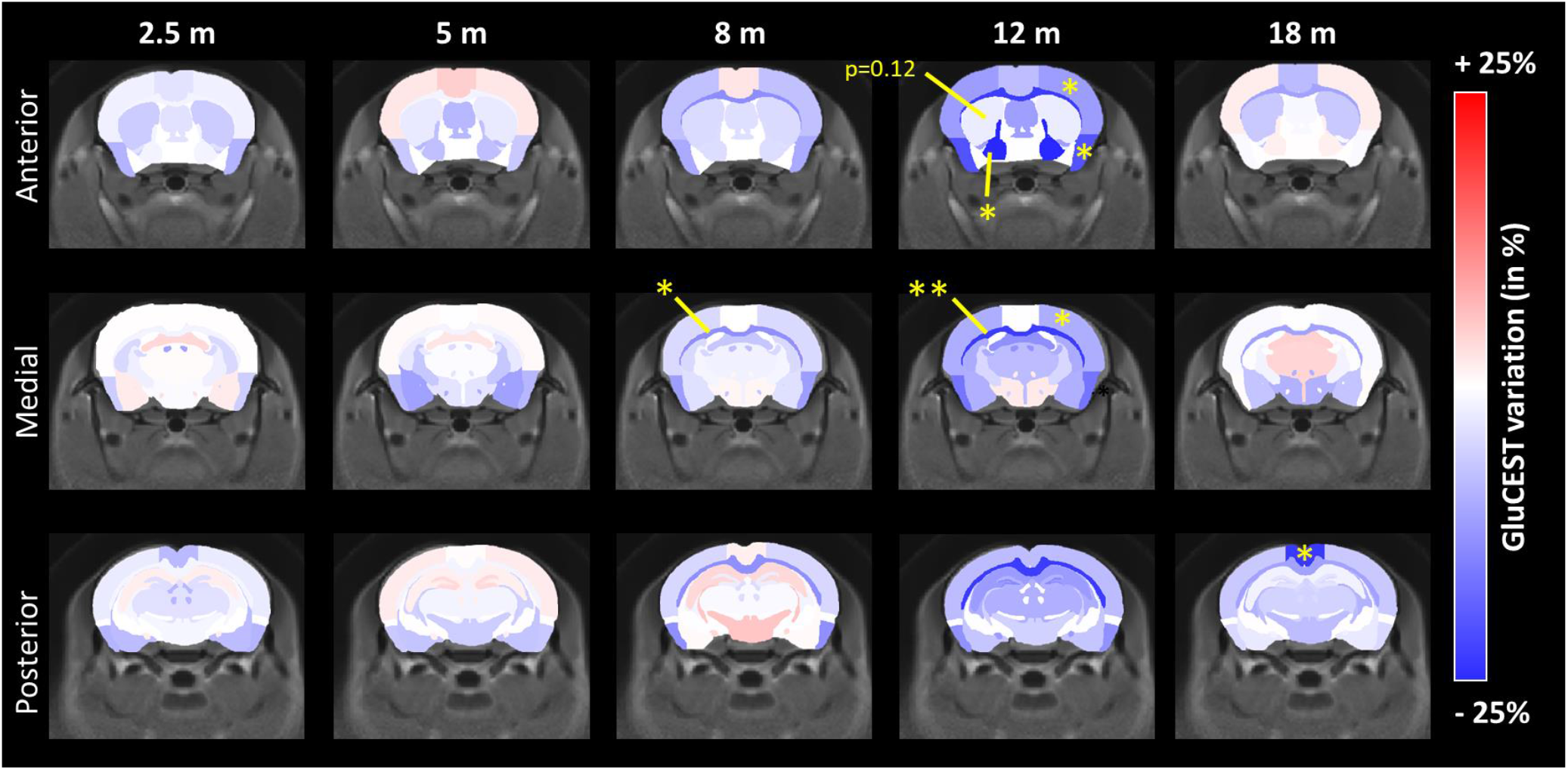
GluCEST variation maps between WT and Ki140CAG mice, at each time-point, on three slices of the mouse brain. Structures with increased gluCEST contrast are reported in shades of red and structures with decreased gluCEST contrast in shades of blue. Yellow stars represent significant difference (RM-ANOVA + Bonferroni, p < 0.05).

Similar maps of MT contrast variations (**Fig. 4**, increase of MT contrast in shades of orange, decrease in shades of purple) revealed significant decrease of MT contrast in the septum (**Fig. 4**, −21.7%, p < 0.05) and a trend in the striatum (−12.5%, p = 0.12) at 12 months only.

**Figure 4:**
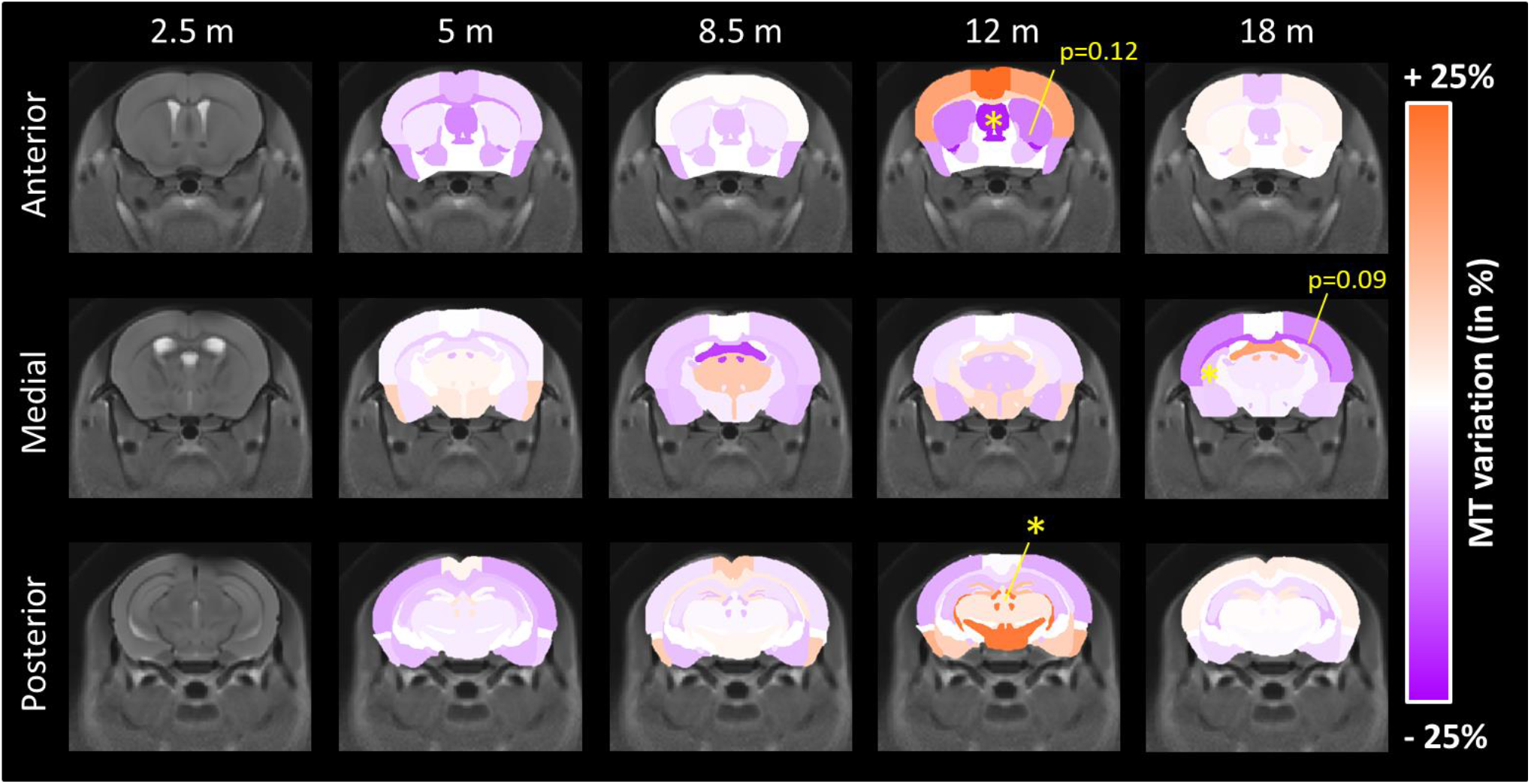
MT variation maps between WT and Ki140CAG mice, at each time-point, on three slices of the mouse brain. Structures with increased MT contrast are reported in shades of orange and structures with decreased MT contrast in shades of violet. Yellow stars represent significant difference (RM-ANOVA + Bonferroni, p < 0.05).

Surprisingly, at 18 months, neither gluCEST nor MT imaging showed significant variation of MTRasym values between WT littermates and Ki140CAG mice. This could be attributed to normal aging of control mice with the decrease of gluCEST and MT contrast for those mice ^65^.

### 3.3 *Alteration of structural connectivity and microstructure in* Ki140CAG mice

Microstructural properties and neuronal fibers integrity were investigated using DTI. Variations of FA, AD and RD between HD and WT mice were calculated and showed for the anterior brain slices (**Fig. 5**, increase of FA, AD or RD in shades of brown, decrease in shades of cyan). Despite a clear tendency of decreased FA, AD and RD values over time in Ki140CAG in most of GM structures, no variation reached statistical significance.

**Figure 5:**
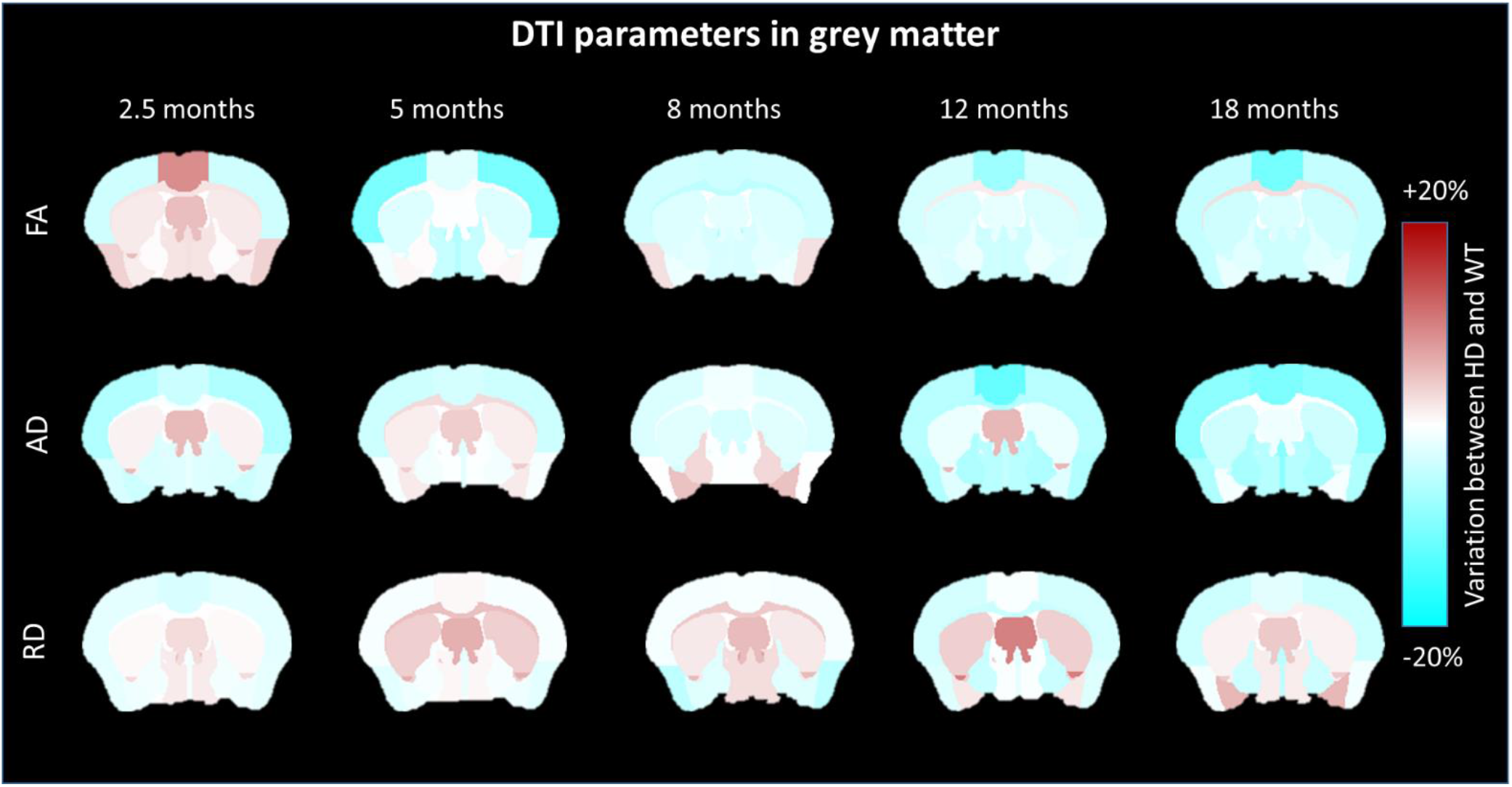
FA, AD and RD variation maps between WT and Ki140CAG mice, at each time-point, on an anterior slice of the mouse brain. Structures with increased diffusion parameters are reported in shades of brown and structures with decreased diffusion parameters in shades of cyan.

We hypothesized that the relevance of atlas-based approach might be limiting for examination of the WM. Indeed, due to small CC thickness and imprecise realignment of WM structures, atlas-based analysis was not able to detect any change. Interestingly, both HD and WT groups showed increase of FA values during the first 8 months and then reached a plateau (**Fig. 6.a**). Nonetheless, both time courses were not identical and a delay of FA increase was observed in Ki140CAG mice. Consequently, we performed TBSS analysis to improve DTI data analysis for WM structures. Thanks to TBSS approach, several clusters of voxels with significant decrease of FA values were identified in the anterior CC of Ki140CAG mice as early as 5 months (**Fig. 6.b**). Clusters of reduced FA values were also observed in the anterior CC at 8 and 12 months in very precise locations confirming the local alteration of the anterior CC that could not be detected with atlas-based approach.

**Figure 6:**
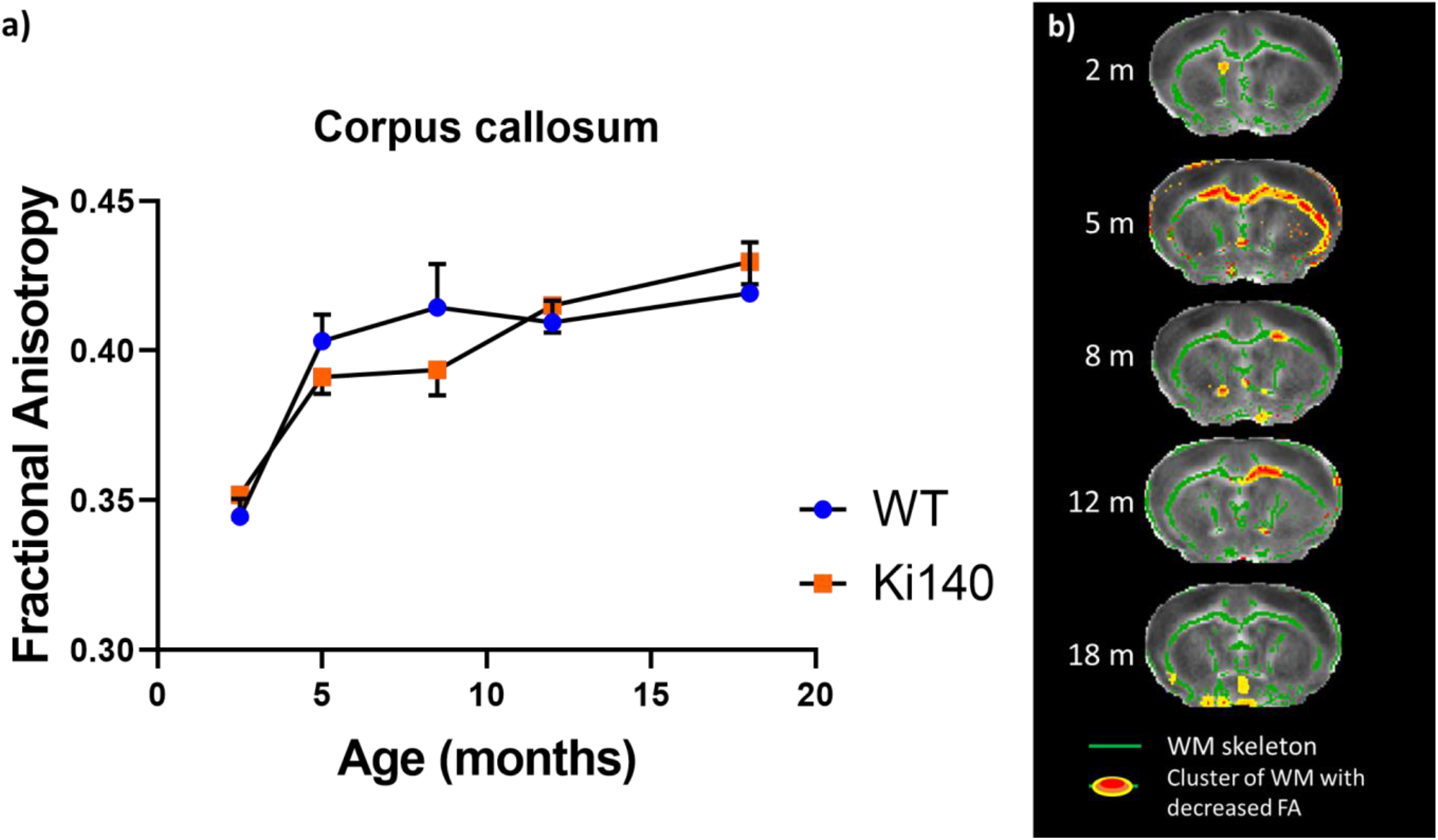
**a.** FA values measured in CC of WT and Ki140CAG mice as a function of time (blue and orange dots respectively, mean ± SEM). **b.** FA alterations in white matter as revealed by TBSS analysis, at each time-point, on an anterior slice of the mouse brain. Green regions represent mean white matter skeleton as used in TBSS pipeline. Red-yellow zones show clusters of voxels with significant decrease of FA (Threshold-free cluster enhancement, p < 0.05).

### 3.4 Integration of affected structures within anatomical networks

In order to further explore the importance of the connections between brain regions, we confronted our results to Allen Connectivity Atlas ^62^. Precisely, the matrix assessing the normalized strength of axonal projections between all the regions of the mouse brain, available from Oh et al. ^62^, was represented using graph theory. **Figure 7** represents physical connections in the mouse brain as derived from Oh et al. ^62^ as a network of structures linked with axonal projections. Position of the regions were determined in an unbiased way by force atlas algorithm. Large structures, similar to those we used in our MRI atlas, were identified in this graph (**Fig. 7**, encircled areas). Interestingly, the Caudate-Putamen (CP) is central in the graph, showing its importance in terms of connections with surrounding structures such as Frontal Cortex, Motor Cortex or Pallidum. In order to summarize findings of this study, we superimposed on this network all structures that were found to be altered in one of the MRI modalities (**Fig. 7**, in blue for gluCEST defects, violet for MT defects and green for atrophy). Colored arrows, representing projection from an affected region, seem to link striatal and cortical regions. This representation merging brain connectivity and altered regions identified using our multimodal MRI approach indicates that damaged/dysfunctional structures/pathways in the HD model belong to a relevant anatomical network.

**Figure 7:**
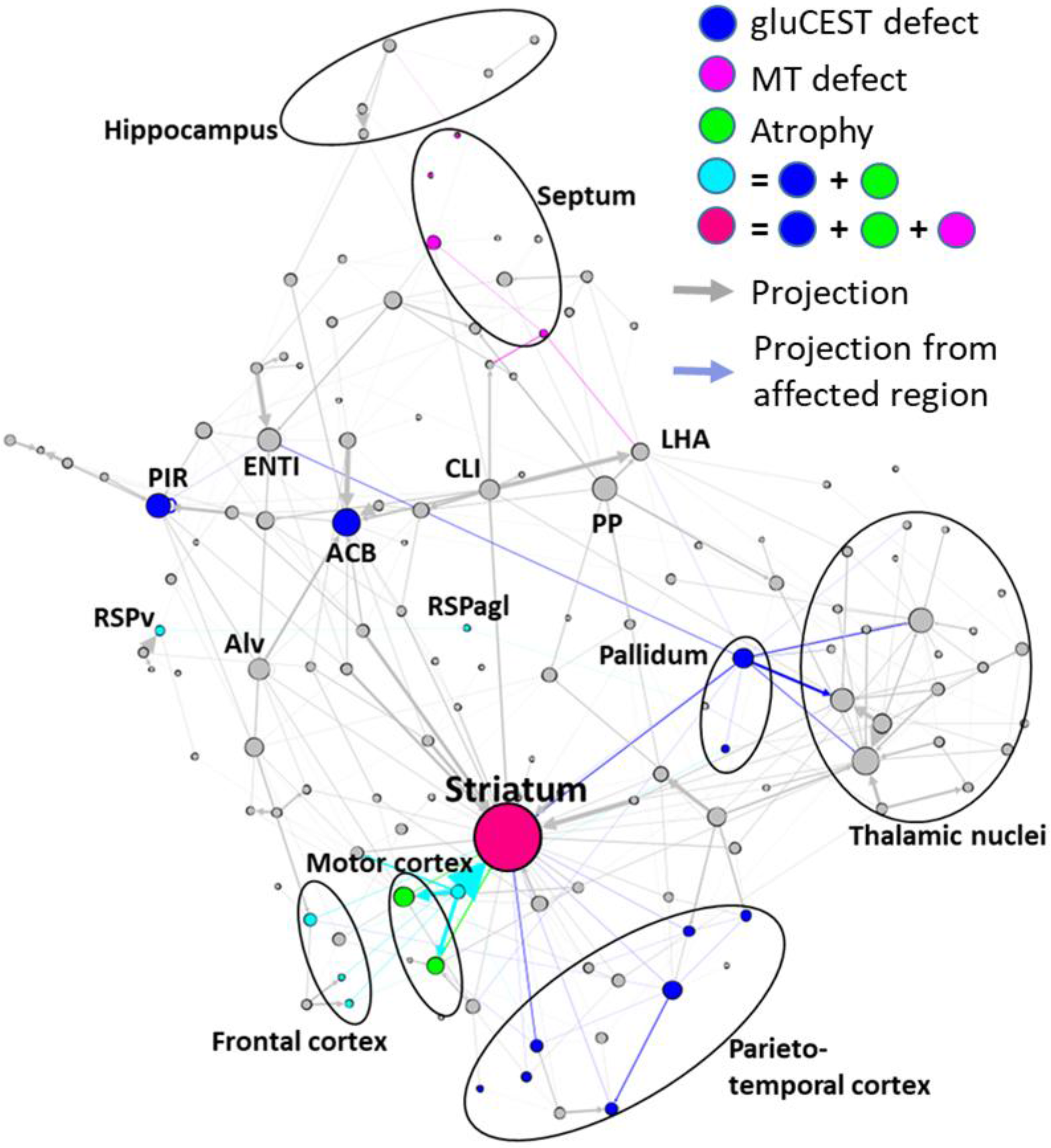
Summary of main results overlaid with graph-theory representation of axonal projection atlas of the mouse brain. Grey arrows and dots represent axonal projections and regions respectively. Colored dots represent affected regions according to our results. Colored arrows represent axonal projections between affected regions and can be associated with white matter alterations. For clarity, only highest degree nodes and affected nodes are labeled. ACB: Nucleus Accumbens; Alv: Alveus; CLI: Central linear nuclei raphe; ENTl: Lateral entorhinal area; LHA: Lateral hypothalamic area; PIR: Piriform area; PP: Peripeduncular nucleus; RSPv: Ventral retrosplenial area; RSPagl: Lateral agranular retrosplenial area.

## 4. Discussion

### 4.1 Ki140CAG as a slowly progressive mouse model of HD

Knock-in models of HD, especially the Ki140CAG model, have been designed and generated through homologous recombination to faithfully replicate the human genetic defect in the mouse HdH gene. This model, along other Knock-in models, is characterized by a mild progression of the disease, as compared to transgenic models where a short fragment of the mutant human HTT gene is inserted randomly in the mouse genome (e.g. R6/2, R6/1 and 171-82Q mice ^54,66^). The Ki140CAG model has already been characterized by slow progression of locomotor performances, behavioral, cellular and molecular abnormalities associated with neurodegeneration ^53^. Moreover, we demonstrated in a previous study that both heterozygous and homozygous Ki140CAG mice exhibit a significant alteration of their metabolic profiles in the striatum as compared to WT littermate controls at 12 months of age.

More interestingly, gluCEST imaging revealed that the CC is the most affected structure in both genotypes, suggesting that this structure is preferentially affected in HD and might play a major role in the physiopathology of the disease ^52^. However, this study only focused on 12-month old mice while numerous studies demonstrated that variations of MRI/MRS indexes or biomarkers are not constant overtime ^67^. Consequently, we thought it was crucial to characterize these changes longitudinally to identify the most relevant biomarkers to monitor disease progression. Indeed, such biomarkers could be of high value to assess the efficacy of experimental therapeutics over time in preclinical trials in genetic rodent models of HD, and if translatable to clinical set ups, they could be a great help in clinical trials.

In the present study, we monitored longitudinally several potential biomarkers to better understand the physiopathology in this particular mouse model of HD. The most commonly used biomarker to characterize HD progression in m-HTT gene carriers is striatal atrophy. In our knock-in model, striatal atrophy is detected only at 18 months (~5% range with approximately 10 animals per group). Based on our results, we could estimate the sample size to compare groups of control and Ki140CAG models assuming a significance level of 5%, a power of 80%, and two-sided tests. Our results showed that 120 animals would be necessary to detect an effect at 12 months.

Similarly, in human studies, the number of premanifest mutant HTT gene carriers or early stage HD patients to be included to observe a significant atrophy of their striatum is high (>30)^68^. Apart from striatal atrophy we found other indexes that seem to be more sensitive with first FA alterations measured as early as 5 months of age in the anterior CC, followed with gluCEST defect at 8 months in the same structure. Our results support the progressivity of the pathogenesis in the Ki140CAG model and highlight the slight progression of the disease in heterozygous mice, making it a great model for the study of early phase of the disease and for the evaluation of new experimental therapeutics with reasonable number of animal per groups.

### 4.2 Identification of molecular biomarkers of HD

From the perspective of identifying relevant and early biological markers of HD, molecules related to energy metabolism and neurotransmission are likely good candidates since it is conceivable that their alteration could occur before actual neurodegeneration associated with major cell shrinkage and/or death of brain cells^69^. Glutamate is an amino acid present in high concentrations in the brain. It is involved in numerous brain functions and plays a major role in brain energy metabolism. [Glu] as measured using ^1^H-MRS has been reported to be affected in the striatum and cortex of HD patients, with conflicting results on whether it increases or decreases ^70,71^. [Glu] decreases in the striatum have also been observed in zQ175 and R6/2 mice ^44,45,72,73^. We found similar reduction in [Glu] in the striatum of Ki140 mice^52,74^. However, due to inherently low spatial resolution of ^1^H-MRS, observations have been limited to striatum and cortex. Here, we could evaluate glutamate changes at the whole brain level using gluCEST imaging. This method is able to capture subtle variations of [Glu] with a high spatial resolution and a rather good specificity. Indeed, previous studies demonstrated that approximately 70% of gluCEST contrast originate from Glu ^49^. In a previous study, we showed that gluCEST imaging is able to map deficiencies linked to glutamate content in 12 month-old Ki140CAG mice ^52^. In particular, it detected reduced contrast in the CC of 12-months old heterozygous mice, with increased severity in age-matched homozygous mice. In the present study, with a new cohort of Ki140 mice, we confirmed the 18.8% decrease of gluCEST contrast measured at 12 months in the CC when compared to wild type littermates confirming the good reproducibility of such measurements. We also show that a significant 10.8% decrease of gluCEST contrast can be observed earlier in the anterior part of the CC at 8 months of age. Noticeably, this reduction in gluCEST signal does not result from a “partial volume” bias, since morphometric measures showed the CC volume was similar in Ki140 mice and controls. Interestingly, following timepoints showed a spreading of gluCEST decrease to surrounding structures, such as the motor and piriform cortices, and subcortical nuclei, including the pallidum and the striatum. We hypothesize that this defect is an early manifestation of alteration of the axonal projections between motor cortex and striatum that pass through the anterior corpus callosum as shown in **Figure 8**. Indeed, **Figure 8.a** shows the cortico-striatal and inter-hemispheric tracts isolated using locally-constrained tractography on a representative dataset from this study. Out of 100,000 total streamlines used in whole brain tractography, 8575 were part of the CC, while 5320 were part of both the CC and the cortico-striatal tract. Thus, we can estimate a proportion of about 5320/8575 ~ 62% of streamlines in the CC going through the cortico-striatal tract. The estimation of streamlines belonging to cortico-striatal tract passing though the CC suggests that this specific tract is not negligible in this area of the CC and alterations may be picked out by our measurements. In addition, **Figure 8.b** supports this idea by showing the projections from Primary Motor Area of motor cortex, using more precise cell tracking technique (Allen Connectivity Atlas) with a good correlation to tractography shown in **Figure 8.a**.

**Figure 8:**
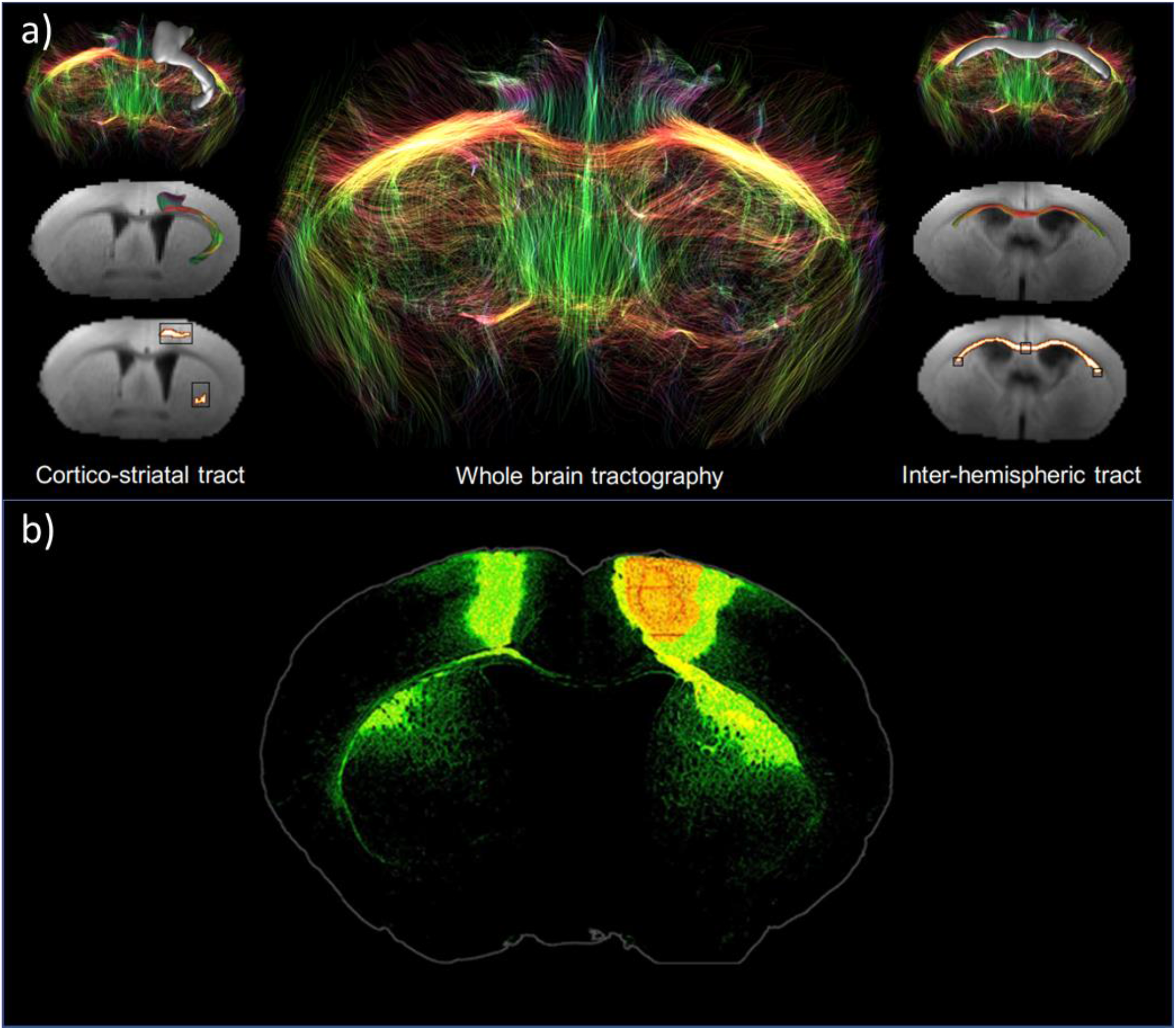
**a.** White Matter tracts isolated from the whole brain tractography of a representative dataset using locally-constrained tractography. Black boxes show starting and ending regions-of-interest (ROIs). White voxels show fibers satisfying this criterion. **b.** Axonal projections from Primary Motor Area. Extracted from Allen Connectivity Atlas (https://connectivity.brain-map.org/projection).

Complementarily to gluCEST, MT is based on the exchange of magnetization between free-water protons and protons of water bound to proteins such as myelin ^36^. Thus, MT signal can be another potential candidate for biomarker identification. Reduced MT contrast can be symptomatic of disruption of myelin sheath or axonal loss ^36,75^. In HD, decreased MT contrast was found in grey matter and white matter of pre-symptomatic gene carriers and HD patients, with correlation between MT loss and disease severity ^76,77^.

In this study, we reported for the first time MT defects in a mouse model of HD, especially in the septum and in the striatum. Such decrease of MT contrast could reflect alteration of myelin contents in affected brain structures as already reported in pre-symptomatic and symptomatic m-HTT gene carriers ^78^.

### 4.3 Altered diffusion in corpus callosum expose the key role of white matter in HD pathogenesis

FA is the most commonly used index to study defects related to microstructural organization. Diffusion of water molecules within WM is highly anisotropic due to strong organization of highly myelinated fibers tracks, providing high FA values. In this study, we observed several clusters of reduced FA values in the anterior part of the CC of Ki140CAG mice at different ages (**Fig. 6.a**). Such results argue in favor of preferential alteration of WM integrity, especially in the anterior part of the CC. Even if interpretation of FA changes is not straightforward^79^, several studies have already showed similar decrease of FA values in the WM of HD patients ^33,80^. Significant alteration of FA and other diffusion parameters were also observed in premanifest HD gene carriers, with a good correlation with WM atrophy ^81^. Similar decrease of FA value was also observed in WM of YAC128 mice as early as 2 weeks of age ^21,22,82^. In our study, we only observed a relatively small part of the CC with reduced FA value. Besides, no variation of CC volume or RD and AD was measured in Ki140CAG but this could be due to limited size of our animal cohorts or to anisotropic voxels used in our study that could bring partial volume effect.

Surprisingly, our results showed a reduction of the difference of FA values between Ki140CAG and WT mice over time (**Fig. 6.b**). Such observation was not consistent with the literature ^21,22,82^. However, one possible explanation is the difference in WM maturation between Ki140CAG mice and their WT littermates. Indeed, development of cortico-striatal tracts was shown affected in Ki111CAG animals ^83,84^ so FA values at early timepoints may be reduced as compared with control animals. Recently, developmental origin of HD has been thoroughly discussed ^83–86^. Cortical progenitor cells from Ki111CAG mice and human gene carriers have shown impaired division in the developmental phase ^83,87^. Compensatory mechanisms may occur in the early life of mice to compensate maturation/developmental defects. However, cortico-striatal projections may remain vulnerable and could explain metabolic impairments observed at 8 months of age.

### 4.4 *Identification of a vulnerable network of brain structures in* Ki140CAG *mice*

In the present study, we performed a longitudinal follow-up of heterozygous Ki140CAG mice with several MRI modalities in order to monitor potential alterations of tissue microstructure, metabolic or energetic defects, and morphological changes of brain structures. Rather than taking these results separately, integrating both structural and functional data may result in a better understanding of the pathogenesis of HD in the Ki140CAG mouse model. We observed that cortical and striatal regions were vulnerable in terms of both atrophy and metabolic changes. In addition, DTI and gluCEST highlighted early WM changes in the anterior part of the CC. Interestingly, striatum and frontal cortex are anatomically connected through the anterior part of the CC and cortico-striatal connections have been shown to be impaired in HD patients as well as in animal models ^88–91^. Our findings corroborate these results and could reflect specific alteration of such connections in the Ki140CAG mouse model.

Thanks to the representation on **Figure 7**, one can observe that striatum seems to be particularly vulnerable as it appears to be affected on several MRI indexes. Moreover, this vulnerability seems to be associated with its centrality in the network and connection strength to cortical tissues that are also affected, through white matter tracts that are early affected.

Based on temporality of the observed modifications and from results found in the literature, we hypothesize that our longitudinal, multiparametric MRI protocol was able to image the vulnerability of the cortico-striato-thalamic glutamatergic pathway^92^, which is known to be impaired in HD. Metabolic and diffusion deficiencies that appear in the white matter might be related to alterations found at the cortico-striatal synapse between Cortical Pyramidal Neurons (CPNs) and Medium Spiny Neurons (MSNs). These alterations include diminished Brain-Derived Neurotrophic Factor (BDNF) transport, increased glutamate neurotransmission, altered NMDA receptors transcription and reduced Ca^2+^ uptake from mitochondria, that together lead to vulnerability of MSNs to excitotoxicity ^93^. Such hypothesis would need further validations with histopathological study to confirm the status of WM tracks at different ages.

## 5. Conclusion

In this study, we developed a multimodal MRI protocol to identify biomarkers of HD pathogenesis. This protocol was applied to monitor longitudinally heterozygous Ki140CAG mice, a slowly progressive mouse model of HD. Thanks to this multimodal approach, we identified early defects in striatal and cortical tissues. Interestingly, MT and gluCEST modalities seemed to be earlier indicator of pathology progression than structure atrophy. Moreover, combination of all results obtained with different MRI modalities were integrated to propose a specific brain network that would be particularly vulnerable in this animal model. As most of these modalities have already been applied on clinical scanners, such approach would be particularly interesting to assess disease progression in a clinical context of HD. This might help identifying more relevant and early biomarkers in premanifest HD patients and could provide relevant information about disease pathogenesis.

## 6. Acknowledgments

This work was supported by two grants from Agence Nationale pour la Recherche (“HDeNERGY” project ANR-14-CE15-0007-01 and “epiHD” project ANR-17-CE12-2019) and one grant from the ERA-Net for Research Programs on Rare Diseases (“TreatPolyQ” project ANR-17-RAR3-0008-01). The 11.7T MRI scanner was funded by a grant from NeurATRIS: A Translational Research Infrastructure for Biotherapies in Neurosciences (“Investissements d’Avenir”, ANR-11-INBS-0011). MP is supported by UKRI Future Leaders Fellowship MR/T020296/1.

## List of Abbreviations

^1^H-MRS: ^1^H Magnetic Resonance Spectroscopy
AD: axial diffusivity
BDNF: Brain-Derived Neurotrophic Factor
CEST: Chemical Exchange Saturation Transfer
Cho: choline
DTI: Diffusion Tensor Imaging
FA: fractional anisotropy
gluCEST: Chemical Exchange Saturation Transfer of glutamate
GM: gray matter
Gln: glutamine Glu: glutamate
HD: Huntington’s disease
HTT: gene coding the huntingtin protein
htt: huntingtin protein
Ins: myo-inositol
MD: mean diffusivity
m-HTT: gene coding the mutant huntingtin protein
m-htt: mutant huntingtin protein
MT: Magnetization transfer
NAA: N-acetyl aspartate
RD: radial diffusivity
RF: radio frequency
TBSS: Tract-Based Spatial Statistics WM: white matter

## Notes

### Competing Interest Statement

The authors have declared no competing interest.

## References

1. Walker, F. O. Huntington’s disease. Lancet Lond. Engl. 369, 218–228 (2007).

2. Liot, G., Valette, J., Pépin, J., Flament, J. & Brouillet, E. Energy defects in Huntington’s disease: Why ‘in vivo’ evidence matters. Biochem. Biophys. Res. Commun. 483, 1084–1095 (2017).

3. Dubinsky, J. M. Towards an Understanding of Energy Impairment in Huntington’s Disease Brain. J. Huntingt. Dis. 6, 267–302 (2017).

4. Paoli, R. A. et al. Neuropsychiatric Burden in Huntington’s Disease. Brain Sci. 7, (2017).

5. Stout, J. C. et al. Neurocognitive signs in prodromal Huntington disease. Neuropsychology 25, 1–14 (2011).

6. Hogarth, P. et al. Interrater agreement in the assessment of motor manifestations of Huntington’s disease. Mov. Disord. 20, 293–297 (2005).

7. Tabrizi, S. J. et al. Potential endpoints for clinical trials in premanifest and early Huntington’s disease in the TRACK-HD study: analysis of 24 month observational data. Lancet Neurol. 11, 42–53 (2012).

8. Bird, E. D. & Iversen, L. L. HUNTINGTON’S CHOREA: POST-MORTEM MEASUREMENT OF GLUTAMIC ACID DECARBOXYLASE, CHOLINE ACETYLTRANSFERASE AND DOPAMINE IN BASAL GANGLIA. Brain 97, 457–472 (1974).

9. Vonsattel, J.-P. et al. Neuropathological Classification of Huntington’s Disease. J. Neuropathol. Exp. Neurol. 44, 559–577 (1985).

10. Kuhl, D. E. et al. Cerebral metabolism and atrophy in Huntington’s disease determined by 18FDG and computed tomographic scan. Ann. Neurol. 12, 425–434 (1982).

11. Brouillet, E., F, C., Mf, B. & P, H. Replicating Huntington’s disease phenotype in experimental animals. Prog. Neurobiol. 59, 427–468 (1999).

12. Tabrizi, S. J. et al. Predictors of phenotypic progression and disease onset in premanifest and early-stage Huntington’s disease in the TRACK-HD study: analysis of 36-month observational data. Lancet Neurol. 12, 637–649 (2013).

13. Aylward, E. H. et al. Onset and rate of striatal atrophy in preclinical Huntington disease. Neurology 63, 66–72 (2004).

14. Paulsen, J. S. et al. Detection of Huntington’s disease decades before diagnosis: the Predict-HD study. J. Neurol. Neurosurg. Psychiatry 79, 874–880 (2008).

15. Kipps, C. M. et al. Progression of structural neuropathology in preclinical Huntington’s disease: a tensor based morphometry study. J. Neurol. Neurosurg. Psychiatry 76, 650–655 (2005).

16. Gómez-Ansón, B. et al. Prefrontal cortex volume reduction on MRI in preclinical Huntington’s disease relates to visuomotor performance and CAG number. Parkinsonism Relat. Disord. 15, 213–219 (2009).

17. Paulsen, J. S. et al. Striatal and white matter predictors of estimated diagnosis for Huntington disease. Brain Res. Bull. 82, 201–207 (2010).

18. Crawford, H. E. et al. Corpus callosal atrophy in premanifest and early Huntington’s disease. J. Huntingt. Dis. 2, 517–526 (2013).

19. Gregory, S. et al. Natural biological variation of white matter microstructure is accentuated in Huntington’s disease. Hum. Brain Mapp. 39, 3516–3527 (2018).

20. Dumas, E. M. et al. Early changes in white matter pathways of the sensorimotor cortex in premanifest Huntington’s disease. Hum. Brain Mapp. 33, 203–212 (2011).

21. Teo, R. T. Y. et al. Structural and molecular myelination deficits occur prior to neuronal loss in the YAC128 and BACHD models of Huntington disease. Hum. Mol. Genet. 25, 2621–2632 (2016).

22. Garcia-Miralles, M. et al. Laquinimod rescues striatal, cortical and white matter pathology and results in modest behavioural improvements in the YAC128 model of Huntington disease. Sci. Rep. 6, 31652 (2016).

23. Huang, B. et al. Mutant huntingtin downregulates myelin regulatory factor-mediated myelin gene expression and affects mature oligodendrocytes. Neuron 85, 1212–1226 (2015).

24. Bartzokis, G. et al. Myelin breakdown and iron changes in Huntington’s disease: pathogenesis and treatment implications. Neurochem. Res. 32, 1655–1664 (2007).

25. Casella, C., Lipp, I., Rosser, A., Jones, D. K. & Metzler–Baddeley, C. A Critical Review of White Matter Changes in Huntington’s Disease. Mov. Disord. 35, 1302–1311 (2020).

26. Zhang, J. et al. In vivo characterization of white matter pathology in premanifest huntington’s disease. Ann. Neurol. 84, 497–504 (2018).

27. Gregory, S. et al. Characterizing White Matter in Huntington’s Disease. Mov. Disord. Clin. Pract. 7, 52–60 (2020).

28. Le Bihan, D. Looking into the functional architecture of the brain with diffusion MRI. Nat. Rev. Neurosci. 4, 469–480 (2003).

29. Sen, P. N. & Basser, P. J. A Model for Diffusion in White Matter in the Brain. Biophys. J. 89, 2927–2938 (2005).

30. Le Bihan, D. & Johansen-Berg, H. Diffusion MRI at 25: exploring brain tissue structure and function. NeuroImage 61, 324–341 (2012).

31. Gregory, S. et al. Longitudinal Diffusion Tensor Imaging Shows Progressive Changes in White Matter in Huntington’s Disease. J. Huntingt. Dis. 4, 333–346 (2015).

32. Rosas, H. D. et al. Complex spatial and temporally defined myelin and axonal degeneration in Huntington disease. NeuroImage Clin. 20, 236–242 (2018).

33. Poudel, G. R. et al. Longitudinal change in white matter microstructure in Huntington’s disease: The IMAGE-HD study. Neurobiol. Dis. 74, 406–412 (2015).

34. Ou, X., Sun, S.-W., Liang, H.-F., Song, S.-K. & Gochberg, D. F. Quantitative Magnetization Transfer Measured Pool Size Ratio Reflects Optic Nerve Myelin Content in ex vivo Mice. Magn. Reson. Med. Off. J. Soc. Magn. Reson. Med. Soc. Magn. Reson. Med. 61, 364–371 (2009).

35. van Zijl, P. C. M., Lam, W. W., Xu, J., Knutsson, L. & Stanisz, G. J. Magnetization Transfer Contrast and Chemical Exchange Saturation Transfer MRI. Features and analysis of the field-dependent saturation spectrum. NeuroImage 168, 222–241 (2018).

36. Tambasco, N. et al. Magnetization transfer MRI in dementia disorders, Huntington’s disease and parkinsonism. J. Neurol. Sci. 353, 1–8 (2015).

37. Bourbon-Teles, J. et al. Myelin Breakdown in Human Huntington’s Disease: Multi-Modal Evidence from Diffusion MRI and Quantitative Magnetization Transfer. Neuroscience 403, 79–92 (2019).

38. Chaney, A., Williams, S. R. & Boutin, H. In vivo molecular imaging of neuroinflammation in Alzheimer’s disease. J. Neurochem. 149, 438–451 (2019).

39. Graham, S. F. et al. Biochemical Profiling of the Brain and Blood Metabolome in a Mouse Model of Prodromal Parkinson’s Disease Reveals Distinct Metabolic Profiles. J. Proteome Res. 17, 2460–2469 (2018).

40. Mochel, F. et al. Early Alterations of Brain Cellular Energy Homeostasis in Huntington Disease Models. J. Biol. Chem. 287, 1361–1370 (2012).

41. Sturrock, A. et al. Magnetic resonance spectroscopy biomarkers in premanifest and early Huntington disease. Neurology 75, 1702–1710 (2010).

42. Zacharoff, L. et al. Cortical metabolites as biomarkers in the R6/2 model of Huntington’s disease. J. Cereb. Blood Flow Metab. 32, 502–514 (2012).

43. Jenkins, B. G. et al. Effects of CAG repeat length, HTT protein length and protein context on cerebral metabolism measured using magnetic resonance spectroscopy in transgenic mouse models of Huntington’s disease. J. Neurochem. 95, 553–562 (2005).

44. Tkac, I., Dubinsky, J. M., Keene, C. D., Gruetter, R. & Low, W. C. Neurochemical changes in Huntington R6/2 mouse striatum detected by in vivo 1H NMR spectroscopy. J. Neurochem. 100, 1397–1406 (2007).

45. Heikkinen, T. et al. Characterization of neurophysiological and behavioral changes, MRI brain volumetry and 1H MRS in zQ175 knock-in mouse model of Huntington’s disease. PloS One 7, e50717 (2012).

46. Wolff, S. D. & Balaban, R. S. Regulation of the predominant renal medullary organic solutes in vivo. Annu. Rev. Physiol. 52, 727–746 (1990).

47. Ward, K. M., Aletras, A. H. & Balaban, R. S. A new class of contrast agents for MRI based on proton chemical exchange dependent saturation transfer (CEST). J. Magn. Reson. San Diego Calif 1997 143, 79–87 (2000).

48. Ward, K. M. & Balaban, R. S. Determination of pH using water protons and chemical exchange dependent saturation transfer (CEST). Magn. Reson. Med. 44, 799–802 (2000).

49. Cai, K. et al. Magnetic Resonance Imaging of Glutamate. Nat. Med. 18, 302–306 (2012).

50. Cai, K. et al. Mapping glutamate in subcortical brain structures using high-resolution GluCEST MRI. NMR Biomed. 26, 1278–1284 (2013).

51. Carrillo-de Sauvage, M.-A. et al. The neuroprotective agent CNTF decreases neuronal metabolites in the rat striatum: an in vivo multimodal magnetic resonance imaging study. J. Cereb. Blood Flow Metab. Off. J. Int. Soc. Cereb. Blood Flow Metab. 35, 917–921 (2015).

52. Pépin, J. et al. In vivo imaging of brain glutamate defects in a knock-in mouse model of Huntington’s disease. NeuroImage 139, 53–64 (2016).

53. Menalled, L. B., Sison, J. D., Dragatsis, I., Zeitlin, S. & Chesselet, M.-F. Time course of early motor and neuropathological anomalies in a knock-in mouse model of Huntington’s disease with 140 CAG repeats. J. Comp. Neurol. 465, 11–26 (2003).

54. Menalled, L. B. & Chesselet, M.-F. Mouse models of Huntington’s disease. Trends Pharmacol. Sci. 23, 32–39 (2002).

55. Hickey, M. A. et al. Extensive early motor and non-motor behavioral deficits are followed by striatal neuronal loss in Knock-in Huntington’s disease mice. Neuroscience 157, 280–295 (2008).

56. Kim, M., Gillen, J., Landman, B. A., Zhou, J. & van Zijl, P. C. M. Water saturation shift referencing (WASSR) for chemical exchange saturation transfer (CEST) experiments. Magn. Reson. Med. 61, 1441–1450 (2009).

57. Celestine, M., Nadkarni, N. A., Garin, C. M., Bougacha, S. & Dhenain, M. Sammba-MRI: A Library for Processing SmAll-MaMmal BrAin MRI Data in Python. Front. Neuroinformatics 14, (2020).

58. Lein, E. S. et al. Genome-wide atlas of gene expression in the adult mouse brain. Nature 445, 168–176 (2007).

59. Liu, G., Gilad, A. A., Bulte, J. W. M., Zijl, P. C. M. van & McMahon, M. T. High-throughput screening of chemical exchange saturation transfer MR contrast agents. Contrast Media Mol. Imaging 5, 162–170 (2010).

60. Smith, S. M. et al. Tract-based spatial statistics: voxelwise analysis of multi-subject diffusion data. NeuroImage 31, 1487–1505 (2006).

61. Smith, S. M. et al. Advances in functional and structural MR image analysis and implementation as FSL. NeuroImage 23, S208–S219 (2004).

62. Oh, S. W. et al. A mesoscale connectome of the mouse brain. Nature 508, 207–214 (2014).

63. Yeh, F.-C., Wedeen, V. J. & Tseng, W.-Y. I. Generalized q-sampling imaging. IEEE Trans. Med. Imaging 29, 1626–1635 (2010).

64. Yeh, F.-C., Verstynen, T. D., Wang, Y., Fernández-Miranda, J. C. & Tseng, W.-Y. I. Deterministic diffusion fiber tracking improved by quantitative anisotropy. PloS One 8, e80713 (2013).

65. Crescenzi, R. et al. Longitudinal imaging reveals sub-hippocampal dynamics in glutamate levels associated with histopathologic events in a mouse model of tauopathy and healthy mice. Hippocampus 27, 285–302 (2017).

66. Menalled, L. B. Knock-in mouse models of Huntington’s disease. NeuroRx J. Am. Soc. Exp. Neurother. 2, 465–470 (2005).

67. Tkac, I. et al. Homeostatic adaptations in brain energy metabolism in mouse models of Huntington disease. J. Cereb. Blood Flow Metab. 32, 1977–1988 (2012).

68. Scahill, R. I. et al. Biological and clinical characteristics of gene carriers far from predicted onset in the Huntington’s disease Young Adult Study (HD-YAS): a cross-sectional analysis. Lancet Neurol. 19, 502–512 (2020).

69. Bonvento, G., Valette, J., Flament, J., Mochel, F. & Brouillet, E. Imaging and spectroscopic approaches to probe brain energy metabolism dysregulation in neurodegenerative diseases. J. Cereb. Blood Flow Metab. 37, 1927–1943 (2017).

70. van den Bogaard, S. J. A. et al. Exploratory 7-Tesla magnetic resonance spectroscopy in Huntington’s disease provides in vivo evidence for impaired energy metabolism. J. Neurol. 258, 2230–2239 (2011).

71. van den Bogaard, S. J. A. et al. Longitudinal metabolite changes in Huntington’s disease during disease onset. J. Huntingt. Dis. 3, 377–386 (2014).

72. Jenkins, B. G. et al. Nonlinear decrease over time in N-acetyl aspartate levels in the absence of neuronal loss and increases in glutamine and glucose in transgenic Huntington’s disease mice. J. Neurochem. 74, 2108–2119 (2000).

73. Peng, Q. et al. Characterization of Behavioral, Neuropathological, Brain Metabolic and Key Molecular Changes in zQ175 Knock-In Mouse Model of Huntington’s Disease. PLOS ONE 11, e0148839 (2016).

74. Pépin, J. et al. Complementarity of gluCEST and 1 H-MRS for the study of mouse models of Huntington’s disease. NMR Biomed. e4301 (2020) doi:10.1002/nbm.4301.

75. Anik, Y., Iseri, P., Demirci, A., Komsuoglu, S. & Inan, N. Magnetization Transfer Ratio in Early Period of Parkinson Disease. Acad. Radiol. 14, 189–192 (2007).

76. van den Bogaard, S., Dumas, E., van der Grond, J., van Buchem, M. & Roos, R. MRI biomarkers in Huntington’s disease. Front. Biosci. Elite Ed. 4, 1910–1925 (2012).

77. van den Bogaard, S. J. A. et al. Magnetization transfer imaging in premanifest and manifest huntington disease: a 2-year follow-up. AJNR Am. J. Neuroradiol. 34, 317–322 (2013).

78. Di Paola, M. et al. MRI measures of corpus callosum iron and myelin in early Huntington’s disease. Hum. Brain Mapp. 35, 3143–3151 (2014).

79. Wheeler-Kingshott, C. A. M. & Cercignani, M. About “axial” and “radial” diffusivities. Magn. Reson. Med. 61, 1255–1260 (2009).

80. Diana Rosas, H. et al. Altered White Matter Microstructure in the Corpus Callosum in Huntington’s Disease: implications for cortical “disconnection”. NeuroImage 49, 2995–3004 (2010).

81. Novak, M. J. U. et al. White matter integrity in premanifest and early Huntington’s disease is related to caudate loss and disease progression. Cortex J. Devoted Study Nerv. Syst. Behav. 52, 98–112 (2014).

82. Petrella, L. I. et al. A whole brain longitudinal study in the YAC128 mouse model of Huntington’s disease shows distinct trajectories of neurochemical, structural connectivity and volumetric changes. Hum. Mol. Genet. 27, 2125–2137 (2018).

83. Barnat, M. et al. Huntington’s disease alters human neurodevelopment. Science 369, 787–793 (2020).

84. Molina-Calavita, M. et al. Mutant Huntingtin Affects Cortical Progenitor Cell Division and Development of the Mouse Neocortex. J. Neurosci. 34, 10034–10040 (2014).

85. Wiatr, K., Szlachcic, W. J., Trzeciak, M., Figlerowicz, M. & Figiel, M. Huntington Disease as a Neurodevelopmental Disorder and Early Signs of the Disease in Stem Cells. Mol. Neurobiol. 55, 3351–3371 (2018).

86. Zhang, C. et al. Abnormal Brain Development in Huntington’ Disease Is Recapitulated in the zQ175 Knock-In Mouse Model. Cereb. Cortex Commun. 1, tgaa044 (2020).

87. Barnat, M., Le Friec, J., Benstaali, C. & Humbert, S. Huntingtin-Mediated Multipolar-Bipolar Transition of Newborn Cortical Neurons Is Critical for Their Postnatal Neuronal Morphology. Neuron 93, 99–114 (2017).

88. Hong, S. L. et al. Dysfunctional Behavioral Modulation of Corticostriatal Communication in the R6/2 Mouse Model of Huntington’s Disease. PLOS ONE 7, e47026 (2012).

89. Naze, S. et al. Cortico-Striatal Cross-Frequency Coupling and Gamma Genesis Disruptions in Huntington’s Disease Mouse and Computational Models. eNeuro 5, (2018).

90. Rebec, G. V. Corticostriatal network dysfunction in Huntington’s disease: Deficits in neural processing, glutamate transport, and ascorbate release. CNS Neurosci. Ther. 24, 281–291 (2018).

91. Blumenstock, S. & Dudanova, I. Cortical and Striatal Circuits in Huntington’s Disease. Front. Neurosci. 14, (2020).

92. Schwartz, T. L., Sachdeva, S. & Stahl, S. M. Glutamate Neurocircuitry: Theoretical Underpinnings in Schizophrenia. Front. Pharmacol. 3, (2012).

93. Fan, M. M. Y. & Raymond, L. A. N-methyl-D-aspartate (NMDA) receptor function and excitotoxicity in Huntington’s disease. Prog. Neurobiol. 81, 272–293 (2007).

